# Increasing TET expression and 5-hydroxymethylcytosine formation by a carbocyclic 5-aza-2’-deoxy-cytidine antimetabolite

**DOI:** 10.1101/2025.11.26.690736

**Authors:** Maike Däther, Elsa Peev, Annika Fröhlich, Binje Vick, Sogol Fatourechi, Gilles Gasparoni, Matthias Heiss, Corinna C. Pleintinger, Emmanuel Asu Bisong, Hans Hurmiz, Davide Guglielminotti, Yasmin V. Gärtner, Tina Aumer, Karsten Spiekermann, Jörn Walter, Irmela Jeremias, Franziska R. Traube, Thomas Carell

## Abstract

Ten-eleven translocation (TET) enzymes are critical epigenetic regulators, which oxidize the methylated cytosine nucleobase 5-methyl-dC (5mdC) in the genome to 5-hydroxymethyl-dC (5hmdC) in an α-ketoglutarate-dependent manner. Because the presence of 5mdC in the promoter region of a given gene silences its expression, this oxidation goes in hand with the reactivation of such silenced genes. In different highly aggressive cancers such as acute myeloid leukemia (AML) and glioblastoma, loss of TET enzyme function and therefore reduced 5hmdC levels pave the way for tumor development. Impairment of TET activity can occur through metabolic inhibition, through loss-of-function mutations in TET genes themselves, and finally through suppression of TET-expression via epigenetic silencing. Reactivation of TET enzyme expression represents a major aim of epigenetic cancer therapy. Here we show that the carbocyclic antimetabolite 5-aza-2’deoxycytidine (cAzadC), which is supposed to suppress the methylation of DNA during replication, leads to a substantial increase of TET2 expression and strongly increasing 5hmdC levels. We show that the treatment with cAzadC goes in hand with the broad reactivation of the cellular anti-tumor responses. With patient-derived xenograft AML-mouse models, we show that this translates into a strongly improved anti-cancer effect *in vivo*.

---

The epigenetic control of transcription is a complex process that involves the reversible modification of histone proteins, e. g. via acetylation and methylation chemistry. In addition, it involves the chemical modification of genomic cytosine (dC) nucleosides, particularly in promoter regions (Fig. 1). ^[1–3]^ On this genomic level, cytosines are methylated to 5-methyl-dC (5mdC) by DNA-methyltransferases (DNMTs), leading to reduced transcription of the corresponding genes.^[4]^ Gene reactivation requires the oxidation of the methyl group in 5mdC to 5-hydroxymethyl-dC (5hmdC).^[5–6]^ This oxidation is achieved by α-ketoglutarate (αKG)-dependent TET dioxygenases.^[7–8]^ In cancer, where tumor suppressor genes are often found silenced, reduced TET activity and consequently reduced 5hmdC level are a hallmark.^[9–12]^

**Figure 1:**
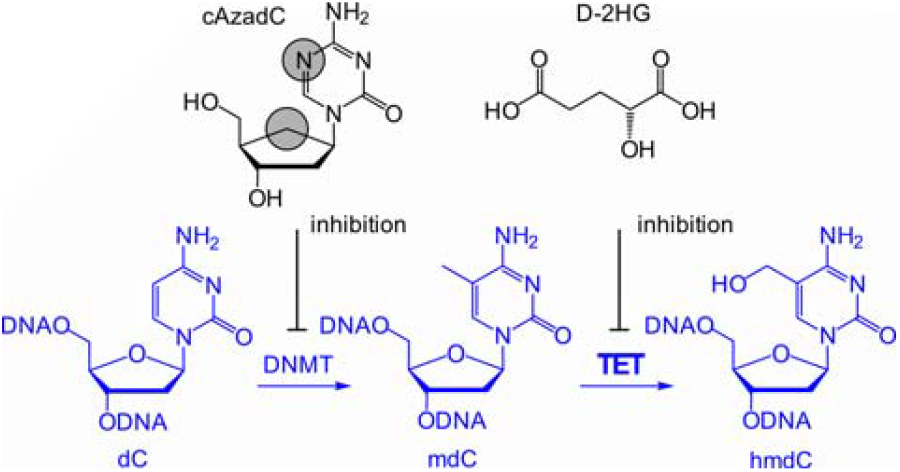
Depiction of the inhibitory cAzadC antimetabolite and the D-2HG onco-metabolite and methylation of dC to 5mdC by DNMT enzymes, followed by the TET-induced oxidation of 5mdC to (5hmdC)

In some tumors, such as many acute myeloid leukemia (AML) and glioma subtypes, neomorphic mutations of isocitrate dehydrogenases 1 and 2 (IDHs), which typically biosynthesize αKG from isocitrate, lead to the production of the onco-metabolite D-2-hydroxyglutarate (2HG), which can accumulate to millimolar concentrations in the tumor cells.^[13]^ At these concentrations, 2HG inhibits the TET enzymes, which prevents the reactivation of genes that would otherwise stop uncontrolled cell growth. ^[14–16]^ Because efficient IDH inhibitors are now available, tumors that feature IDH mutations have a better prognosis.^[17–22]^

In many tumors, however, TET enzymes are epigenetically silenced^[23–24]^ and in these cases, IDH-inhibitors are not expected to improve therapy. A rather unexplored approach to tackle this problem is to increase the expression of TET enzymes.^[25]^ Here we show that this is indeed possible with the carbocyclic 5-aza-2’deoxy-cytidine antimetabolite cAzadC compound shown in Fig. 1.^[26–27]^

We began the investigation by analyzing how well cAzadC would integrate as an antimetabolite into the genome of treated cells upon cell division. Once integrated, we knew that the compound would covalently inhibit the DNMT enzymes via the mechanism shown in Fig. 2A. The result is an inhibition of the DNMT enzymes which leads to a global reduction of the genomic methylation level, as previously shown by us.^[27]^ We investigated the ability of cAzadC to reduce 5mdC levels in different AML cell lines, treating Kasumi-1, HL-60 and for direct comparison again MOLM-13, as well as patient-derived xenograft (PDX) AML cells (AML-491^[28]^) for 72 h with cAzadC at concentrations of 0.5 and 1.0 µM. We used our stable isotope standard based UHPLC-QQQ-MS quantification method^[29]^ in order to measure the levels of cAzadC in the soluble metabolite pool and also in the genome. In addition, we quantified the levels of the epigenetic bases 5mdC and 5hmdC. For the experiment, we first isolated the nucleosides from the soluble cell fraction and we also isolated total DNA which was digested to nucleosides with a nuclease enzyme mixture. We subsequently prepared a mix of synthetic, stable isotope labelled compounds cAzadC*, 5mdC* and 5hmdC* for co-injection into the UHPLC-QQQ-MS system. For quantification, we determined calibration curves. This procedure allowed us to obtain exact quantitative data from the MS-signal intensities. The obtained data are depicted in Figure 2.

**Figure 2:**
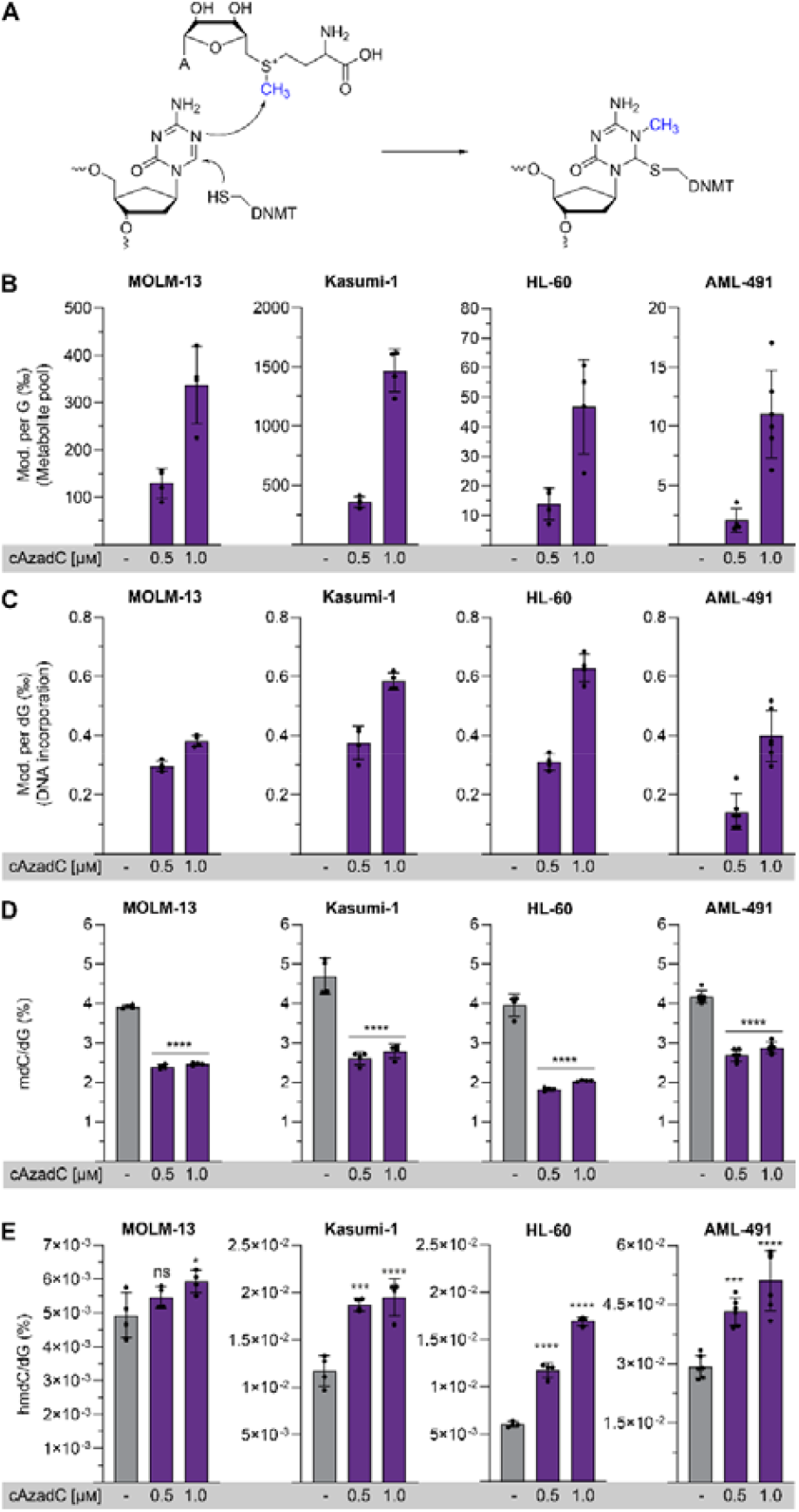
Integration, incorporation and mode of inhibition by cAzadC. (A) Depiction of how cAzadC covalently inhibits DNMT enzymes. (B – E) Quantitative data about the amount of cAzadC in the soluble cell fraction (B) and after genomic incorporation (C) and how treatment with cAzadC affects the levels of 5mdC (D) and 5hmdC (E). All data above are from cells harvested after 72 hours of incubation with DMSO control, 0.5 or 1 µM. The samples were provided in at least 4 biological replicates. The samples were proceeded as described in the methods and analysed by UHPLC-QQQ-MS. The quantitative values were determine using heavy labelled ISTD and then normalized per hundred or thousand dG. Each dot represents the result from one biologically independent replicate. Statistical analysis was performed using two-sided t-test. **** p-value < 0.0001, *** p-value < 0.001, ** p-value < 0.01, * p-value < 0.05, ns = not significant.

Based on literature data, we expected that replacement of the ribose by a cyclopentane unit would negatively affect the ability of cAzadC to be integrated into the genome.^[30]^ This is indeed observed. While the soluble pool contained significant amounts of cAzadC even after 72 h (Fig. 2B), the amount of genome-integrated material was very low, although clearly detectable (Fig. 2C). In the AML cells lines and in the PDX AML-491 cells, we observed an incorporation level of about 0.5 ‰ per dG at 1 μM dosing, which corresponds to the presence of about one million cAzadC molecules in the genome. This is very low compared to other anti-metabolites, but enough to see an improved cell death after a cellular division (72 h), as presented on FACS data (Fig. S1C). As previously shown, the integration is sufficient to deplete the entire DNMT1 pool.^[31–32]^

Despite the low integration levels, we saw in all cell lines, and particularly in the PDX AML-491 cells, a significant reduction of the 5mdC levels to about 35% (AML-491) and 55% (HL-60) (Fig. 2D), showing that the low level of integration does not reduce the epigenetic effect.

In contrast, when we measured 5hmdC, we detected substantially increased levels, ranging between 15% (MOLM-13) and 300% (HL-60) increase (Fig. 2E). Importantly, also in the patient-derived AML-491 leukemic cells, the 5hmdC levels increased by 66% from 0.03% to 0.05% relative to dG. This surprising observation might be best explained by profound changes in gene expression, potentially caused by global genomic demethylation of the genome, but clearly requires further investigation.

Among the other genes monitored, it is noteworthy that growth-inhibiting p53 signaling was highly upregulated, while cancer-promoting MYC signaling was highly downregulated (Fig. 3C, Fig. S1A) which is in line with the idea that the reduction of methylation levels creates a strong anti-proliferative response (Fig. S1B). Interesting and potentially clinically relevant is the observation that the anti-apoptotic protein BCL-2, which is often overexpressed in leukemic cells and in the clinic, is inhibited by treatment with Venetoclax^[38]^, was significantly downregulated after cAzadC treatment (Fig. 3A). In line with these observed transcriptomic changes, cAzadC-treated MOLM-13 cells underwent apoptosis (Fig. S1C).

**Figure 3:**
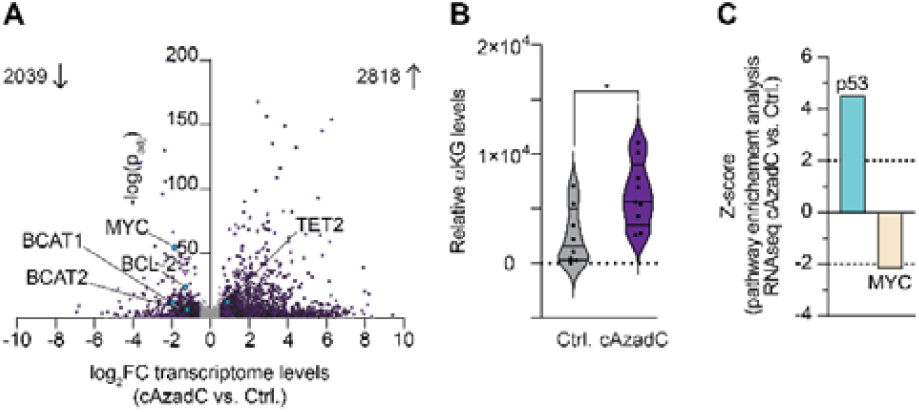
Addressing the hallmarks of cancer in MOLM-13 cells with cAzadC treatment. (A – C) Cells were treated as indicated once with 0.5 µM of cAzadC for 72 h and the effects were compared to DMSO-treated Ctrl. (**A**) Volcano plot of transcriptomic changes. RNAseq analysis with n = 3 biologically independent replicates per condition. The significance threshold for differential expression was set to −log(p_adj_) > 1.3 and |log_2_FC| ≥ 0.5849 (|fold change| ≥ 1.5). treatments. (**B**) Relative αKG levels determined by a fluorescence assay after treatment. Violin plot (including 25th percentile (lower border), median (middle border) and 75th percentile (upper border)) with each dot representing the result from one biologically independent replicate. Statistical analysis was performed using two-sided t-test. * p-value_j_ < 0.05, ns = not significant. (**C**) Results of the pathway enrichment analysis using Ingenuity Pathway Analysis^[39]^.

To investigate how patient-derived AML-491 cells would react, we first quantified the levels of TET2 mRNA after treatment. In these studies, we included the clinically established AzadC compound (Decitabine)^[40–41]^ as a reference. While the Decitabine treatment yielded highly inconsistent TET2 expression changes, the cAzadC compound increased the TET2 expression robustly in a dose-dependent fashion more than 20-fold at the highest concentration of 3 µM (Fig. 4A). This is a dramatic effect and can only be explained by assuming that the expression of the epigenetically acting enzyme TET2 is indeed tightly repressed by DNA methylation and that the global demethylation, caused by cAzadC, leads to an unleashing of its expression. Whatever the exact reason may be, the data show that the complex genetic rewiring processes that are associated with global epigenetic demethylation caused by cAzadC leads to a substantial increase of the amount of TET2. This, in combination with higher αKG levels is suggested to be responsible for the increasing 5hmdC levels.

**Figure 4:**
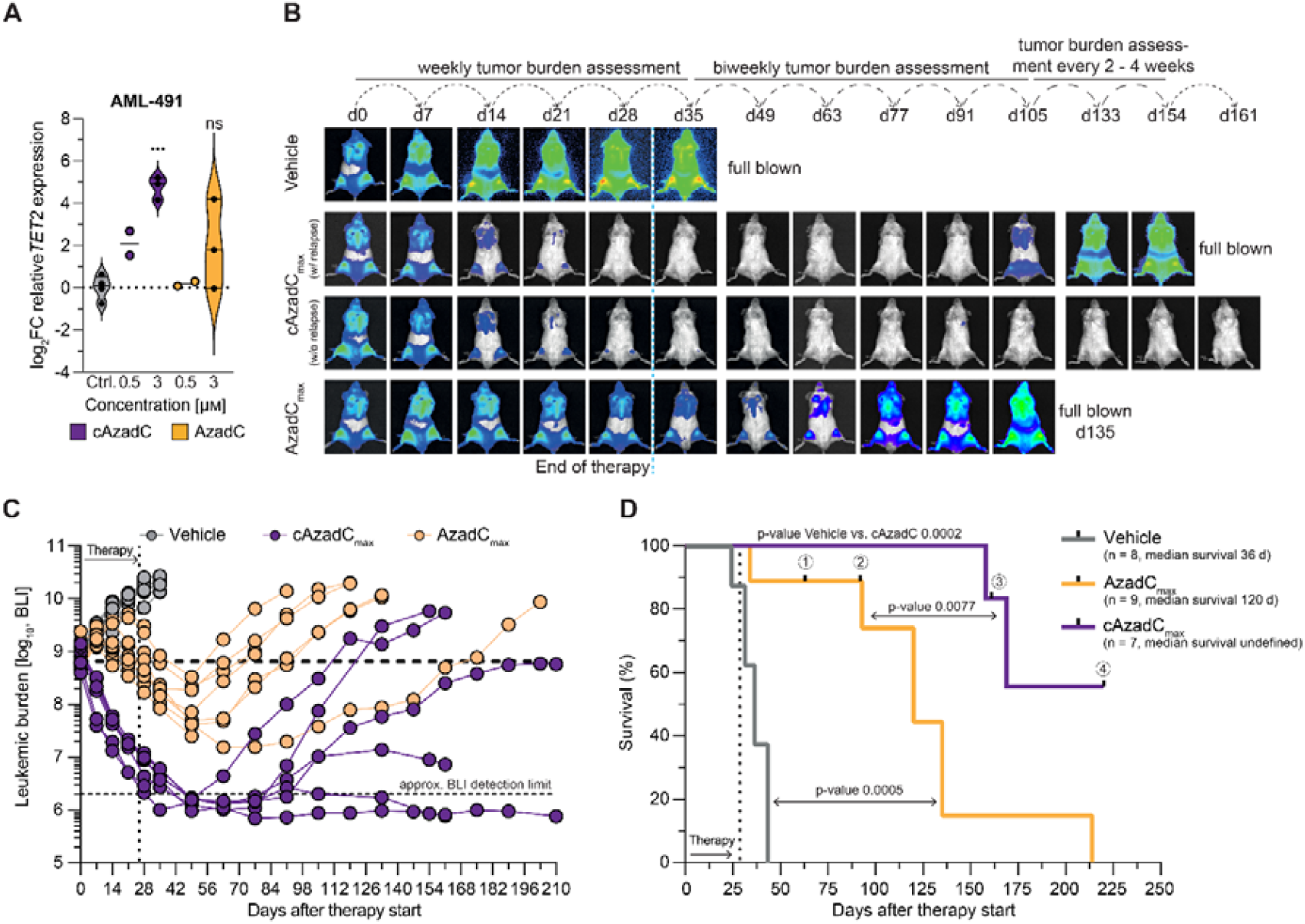
Effects of cAzadC treatment compared to AzadC treatment and vehicle *in vivo*. (A) log_2_ fold-change of TET2 expression levels in AML-491 cells after *ex vivo* treatment with cAzadC compared to AzadC. (**B** – **D**) NSG mice were transplanted iv with luc-positive AML-491 PDX cells. Leukemic burden was monitored by repeated bioluminescence in vivo imaging (BLI). At total flux of 7 × 10^8^ Photons/second, mice were treated with 40 mg/kg cAzadC (cAzadCmax) or 0.25 mg/kg AzadC (AzadCmax) or vehicle as control i.p. for 5 days a week for four weeks. Representative BLI images (**B**) and quantification (**C**), where each curve represents one mouse. (**D**) Kaplan-Meier curves of mice treated with vehicle (n = 8 of 3 independent experiments), AzadCmax (n = 9 of 3 independent experiments) and cAzadCmax-treated (n = 7 of 2 independent experiments). Median survival after therapy start: vehicle (grey): 36 d, AzadCmax (orange): 120 d, cAzadC (violet): undefined. Log-rank (Mantel-Cox) test.

Because increasing 5hmdC levels and higher TET2 activities were in the past firmly associated with reduced tumor growth^[42–43]^, we reasoned that in comparison to Decitabine, cAzadC would show improved antitumor activities.

In order to investigate this, we performed an *in vivo* treatment experiment. We injected luc+ AML-491 PDX cells into NSG recipient mice, which are immunodeficient to accept engraftment of primary human cells, and measured tumor outgrowth by following *in vivo* bioluminescence (BLI). 21 days after injection (referred to as d0), mice were treated for four weeks with either cAzadC, Decitabine or vehicle. Knowing about the different genomic integration kinetics of cAzadC and Decitabine, we decided to perform the comparison at the respective maximally tolerated doses determined previously in a toxicologic study. There, an over 100fold better tolerance was observed for cAzadC compared to Decitabine. While this limited us to 0.25 mg/kg per day over four weeks for Decitabine, we were able to increase the dose to 40 mg/kg per day for cAzadC. The comparison at these maximum tolerated doses showed that, consistent with our speculation, a more rapid decrease in leukemic burden is observed with cAzadC (Fig. 4B, C). The tumor reduction also lasted longer, for even up to three weeks after the end of treatment (Fig. 4B, C). Interestingly, the re-growth of tumor cells at the end of therapy was greatly reduced in both Decitabine and cAzadC-treated mice compared to AML-491 cells of the control that have not undergone treatment, but more pronounced for cAzadC (Fig. 4C, Fig. S2). In two mice, leukemic stem cells (LSCs), expected to be the source of relapse, remained functional as indicated by the resumption of leukemic cell proliferation. In one mouse, leukemic cell proliferation resumed, but stopped before reaching again the BLI values at the beginning of the therapy. In another mouse, proliferation initially resumed, but then stopped and dropped without further treatment below the BLI detection limit, suggesting exhaustion of leukemic cells. Remarkably, in two mice, the leukemic burden remained below the detection limit until they had to be sacrificed due to leukemia-unrelated end-points, indicating that LSCs were effectively eradicated (Fig. 4C). In contrast, Decitabine treatment at the maximum tolerated dose failed to eradicate LSCs, leading in all mice to resumption of proliferation. Despite a significantly increased survival time, all Decitabine-treated mice had to be sacrificed prematurely due to high leukemic burden. Overall, we observed a significant increase in survival of cAzadC treated mice, not only in comparison to control-treated mice, but also compared to mice treated with Decitabine at the maximum tolerated dose (Fig. 4D).

## DATA AVAILABILITY

The RNAseq data of 0.5 µm cAzadC or AzadC treated MOLM-13 cells were deposited at the Gene Expression Omnibus (GEO) repository ^[44–45]^ with the dataset identifier GSE225154.

## Acknowledgements

We thank the Deutsche Forschungsgemeinschaft (DFG) for financial support for this project via CRC1309 (Grant Nr. 325871075, Projects A04 (TC), A05 (JW) and C08 (FRT)), CRC1361 (Grant Nr. 393547839, Project 2 (TC)), TRR237 (Grant Nr. 369799452, Project A27 (TC)). Further support is acknowledged from the BMBF in the framework of the Zukunftscluster program (Cluster for Nucleic Acid Therapeutics Munich, CNATM) (Project ID: 03ZU1201AA (FRT, IJ and TC)). FRT thanks the Daimler und Benz Stiftung (Grant Nr. 32-09/21) and the Fonds der Chemischen Industrie (Liebig Fellowship Li 210/06) for support. This project has received funding from the European Research Council (ERC) under the European Union’s Horizon 2020 research and innovation program under grant agreement Nr. 741912 (EPiR) (TC), the Marie Sklodowska-Curie grant agreements Nr. 861381 (EP) and Nr. 101072780 (DG). MD thanks the Fonds der Chemischen Industrie for a PhD fellowship. YVG was supported by the Bavarian Graduate program RNAmed. IJ thanks German Cancer Aid for support via a Mildred Scheel Professorship. The authors thank Markus Müller for critical discussions and help with manuscript preparation.

## Notes

### Competing Interest Statement

The authors have declared no competing interest.

### Summary of Updates

The acknowledgement section was revised to accommodate MSCA action no. 101072780 (ISOBIOTICS), which was missing in the original submission.

