## Supplementary Material for "Increasing TET expression and 5-hydroxymethylcytosine formation by a carbocyclic 5-aza-2’-deoxy-cytidine antimetabolite"

[1] M. Sc. M. Däther, M. Sc. E. Peev, M. Sc. S. Fatourechi, Dr. M. Heiss, Dr. C. C. Pleintinger, M. Sc. E. A. Bisong, M. Sc. H. Hurmiz, M. Sc. D. Guglielminotti, M Sc. Y. V. Gärtner, Dipl. Pharm. T. Aumer, Jun.- Prof. Dr. F. Traube, Prof. Dr. T. Carell
Institute for Chemical Epigenetics, and Center for Nucleic Acid Therapies, Department of Chemistry, LMU Munich, Butenandtstr. 5 – 13, 81377 Munich, Germany

[2] A. Fröhlich, Dr. B. Vick, Prof. Dr. I. Jeremias
Research Unit Apoptosis in Hematopoietic Stem Cells, Helmholtz Munich, German Research Center for Environmental Health (HMGU), Feodor-Lynen-Str. 21, 81377 Munich, Germany

[3] Dr. G. Gasparoni, Prof. Dr. J. Walter
Department of Genetics/Epigenetics, Universität des Saarlandes, 66041 Saarbrücken, Germany

[4] Prof. Dr. K. Spiekermann, Medizinische Klinik und Poliklinik III, Campus Großhadern, LMU University Hospital, LMU Munich, Marchioninistr. 15, 81377 Munich, Germany

[5] Jun.- Prof. Dr. F. Traube, Institute of Biochemistry, University of Stuttgart, Allmandring 31, 70569 Stuttgart, Germany

[6] Dr. B. Vick, Prof. Dr. I. Jeremias: German Cancer Consortium (DKTK), partner site Munich, a partnership between DKFZ and University Hospital LMU Munich, Munich, Germany

### These authors contributed equally.

#### Supporting Figures

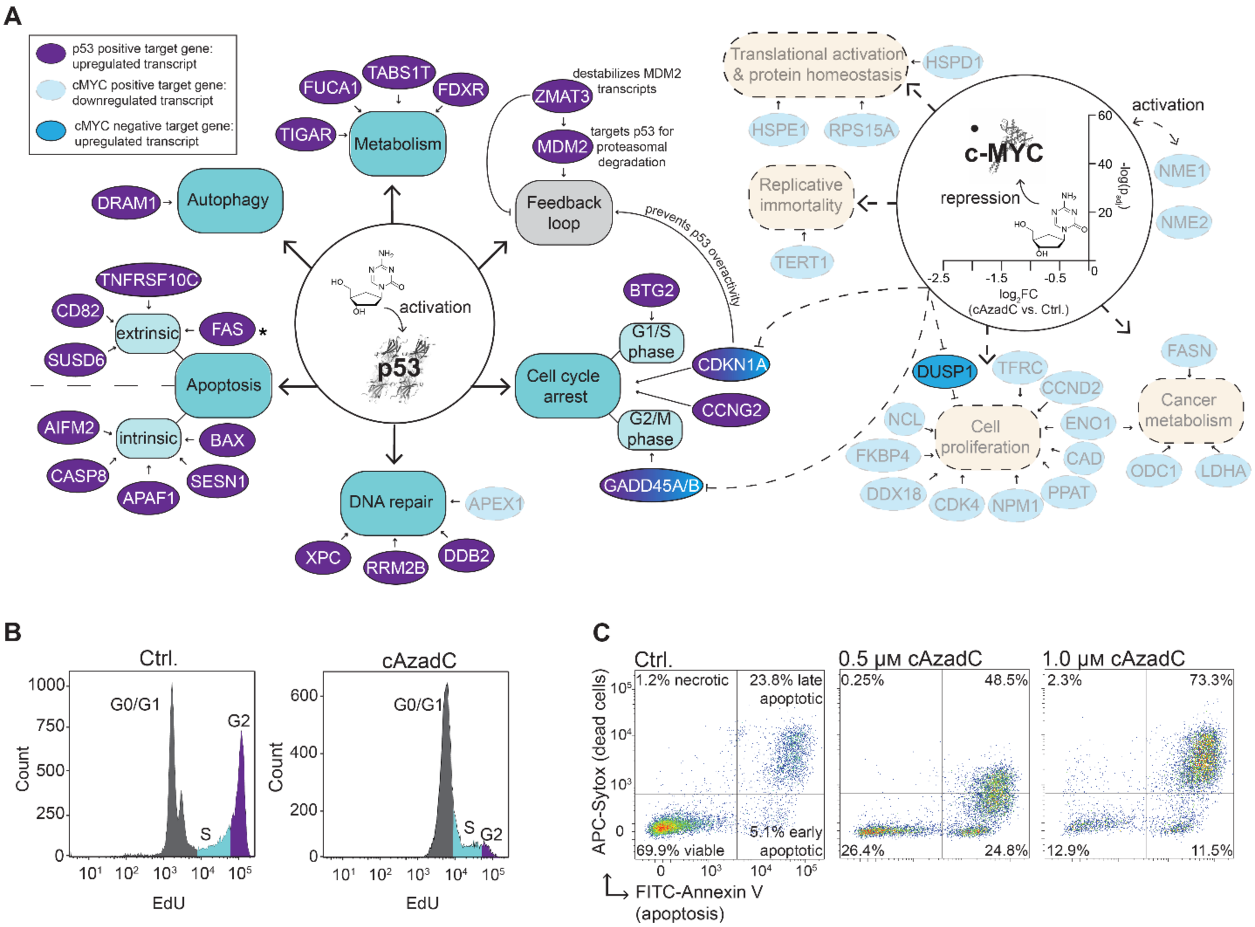


Figure S1: **Anti-leukemic effect of cAzadC**. MOLM-13 cells were treated for 72 h with 0.5 µM (**A** – **C**) and 1.0 µM (**C**) cAzadC. (**A**) Transcription level changes of p53 and MYC-target genes after treatment based on the RNAseq data presented in Fig. 3. Solid lines indicate upregulated target genes; dashed lines indicate downregulated target genes. The genes are grouped according to the cellular process, in which they are involved. (**B**) Proliferation as indicated by EdU incorporation, followed by Click chemistry using an azide-tagged fluorophore and quantification by flow cytometry. (**C**) Apoptotic cells quantified by a flow-cytometry assay based on Annexin V signal intensity. (B – C) 10000 events were quantified, representative result of one out of three independent experiments.

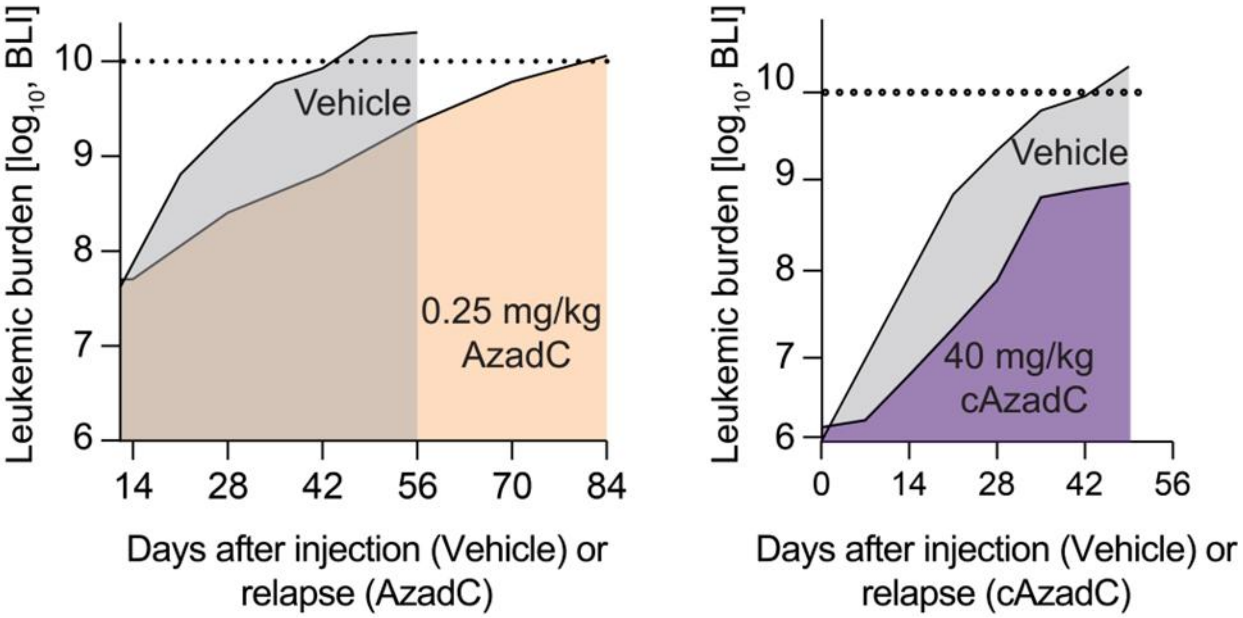


Figure S2: **Growth curves (mean) of relapsed AML-491 cells after treatment with AzadC (0.25 mg/kg) or cAzadC (40 mg/kg) in comparison to growth of treatment-naïve AML-491 cells (Vehicle).**

**Materials and Methods**

General cell culture MOLM-13, Kasumi-1 and HL60 cells

MOLM-13 cells (Leibniz Institute DSMZ – German Collection of Microorganisms and Cell Cultures; AML cell line [1], Kasumi-1 cells (Leibniz Institute DSMZ – German Collection of Microorganisms and Cell Cultures; AML cell line [2] and HL-60 cells (Leibniz Institute DSMZ – German Collection of Microorganisms and Cell Cultures; AML cell line [3]) were cultivated at 37 °C in water-saturated, $CO_2$-enriched (5 %) atmosphere. RPMI-1640 (Merck R0883), containing 20 % (v/v) fetal bovine serum (FBS) (Invitrogen 10500-064) and 2 mm l-alanyl-l-glutamine (Merck G8541) were used as growing medium. When reaching a density of $2 \times 10^6$ cells/mL, the cells were routinely passaged to a density of $0.5 \times 10^6$ cells/mL. Cells were maintained for ~ 25 passages after thawing. Cells were tested regularly (at least once for the time of passaging) for Mycoplasma contamination using Mycoplasma Detection Kit (Jena Bioscience PP-401L).

Cell culture for nucleoside quantification, RNAseq, proliferation assay and apoptosis assay

Cells were seeded at a concentration of $0.5 \times 10^6$ cells/mL unless stated otherwise and directly treated with cAzadC using the indicated concentration and incubation time during which the medium was not renewed. DMSO-treated cells (0.03%) served as a negative control in all experiments unless indicated otherwise.

Purification of DNA and metabolite pool (MP)

For the purification of DNA and MP, cells were washed with DPBS. If cells were not used directly, they were lysed in 800 µL RLT lysis buffer (Qiagen 79216) supplemented with 1% (v/v) 2-mercaptoethanol and stored at -80 °C. gDNA isolation and subsequent digestion for LC-MS/QQQ measurements was performed as previously described [4]. For MP extraction, cells were harvested in 80 % Acetonitrile. The samples were incubated 30 min on ice and occasionally vortexed and then centrifugated 30 min at 4 °C at 10,000 ×g. The supernatant containing the metabolite pool was collected and lyophilized. The dry pellet was then resuspended in buffer containing 50 mm ammonium citrate tribasic and 0.01 % formic acid in LC-MS grade water. Both, digested DNA and MP were filtered through 0.2 µm Supor Natural PP filters (Pall Corporation, AcroPrep Advance 96 Well) before measurement by LC-MS/QQQ.

RNA isolation

400 µL of the first flow-through of the gDNA analysis was used to isolate RNA. If RNA isolation was not performed in parallel to gDNA isolation, the flow-through containing the RNA was 300 µL of 95% Ethanol were added, transferred to a Zymo-Spin IIC column (ZymoResearch C1011-50) and incubated for 1 min. After RNA binding, the column was centrifuged at RT, 1500 *x g* (2 min) and at 10 000 *x g* (30 s). The flow-through was discarded and the column was washed with 800 µL RNA Wash Buffer (ZymoResearch R1003-3). To remove residual DNA contamination, DNA was

digested according to the peqGOLD DNase I Digest kit (VWR 13-1091-01), followed by washing steps with 400 µL RNA Prep Buffer (ZymoResearch R1060-2) and 800 µL RNA Wash Buffer. The RNA was eluted using 53 µL of Milli-Q $H_2O$ and the concentration was determined using a spectrophotometer (Implen, NanoPhotometer N60) in the RNA 40 mode with air bubble recognition set on and 1 µL of sample.

RNA quality and integrity check

After RNA isolation, no further DNA contamination assessment was done. RNA integrity check was done using the Agilent 4150 (G2992AA) TapeStation and Agilent High Sensitivity RNA ScreenTape according to the provided manual for all samples, except for one out of three Ctrl. samples, one additional PBS Ctrl. sample and one out of two 0.5 µM AzadC samples, which were spared because the RNA concentration was not sufficient to run the test. Obtained $RIN^e$ value per independent biological sample:

| Sample | $RIN^e$ |
|---|---|
| AML-491_DMSO_1 | 8.8 |
| AML-491_DMSO_2 | 8.1 |
| AML-491_0.5µM_AzadC_2 | 8.4 |
| AML-491_3.0µM_AzadC_1 | 8.8 |
| AML-491_3.0µM_AzadC_2 | 9.0 |
| AML-491_3.0µM_AzadC_3 | 7.5 |
| AML-491_0.5µM_cAzadC_1 | 8.2 |
| AML-491_0.5µM_cAzadC_2 | 8.5 |
| AML-491_3.0µM_cAzadC_1 | 8.3 |
| AML-491_3.0µM_cAzadC_2 | 8.9 |
| AML-491_3.0µM_cAzadC_3 | 7.7 |

cDNA synthesis and RT-qPCR

cDNA synthesis was performed with the iScript cDNA Synthesis Kit (Bio-Rad 1708891) according to the manufacturer’s protocol using 0.5 - 1 µg of RNA per sample.

For subsequent RT-qPCR the following oligonucleotides were used (HK = house keeper):

| Target | Primer | Sequence (5' – 3') | Tm [°C] | Product length | Start on Target |
|---|---|---|---|---|---|
| ACTB (NM_001101.5) | hActB_fw (HK) | GCCGCCAGCTCACCAT | 59.71 | 119 bp | 71 (exon 2) |
| | hActB_rev (HK) | CACGATGGAGGGGAAGACG | 59.86 | | 189 (exon 2) |
| TET2 (NM_001127208.3) | hTET2_fw | AAGGCTGAGGGACGAGAACGA | 63.22 | 115 bp | 83 (exon 1) |
| | hTET2_rev | TGAGCCCATCTCCTGCTTCCA | 63.29 | | 197 (exon 2) |

Each primer pair (forward and reverse) was mixed and diluted to a final concentration of 1 µM. Subsequently, per 8 µL of diluted primers, 10 µL of iTaq Universal SYBR Green supermix (Bio-Rad 1725124) was added (RT-qPCR reaction mix). RT-qPCR was performed in 20 µL reactions with 2 µL of 10 ng/µL cDNA (20 ng per reaction) mixed with 18 µL of RT-qPCR reaction mix (final primer concentration 400 nM). First, a no-reverse transcription control qPCR was performed for each sample using the hActB primer pair to ensure that no potential DNA contamination would interfere with the experiment. In addition, the hTET2 primer pair was designed to target different exons.

All samples were run in technical triplicates using the following PCR program on a qTOWER$^3$/G cycler (Jena Biosciences):

| | | |
|---|---|---|
| 95 °C | 03:00 min. | |
| 95 °C | 00:10 min. | 40 x |
| 60 °C | 00:20 min. | 40 x |
| 72 °C | 00:30 min. | 40 x |
| Melt | 00:15 min. | |

Using the qPCRsoft 4.1 software, auto threshold mode was selected to calculate the $C_t$ value and the average $C_t$ value for each sample/primer pair was calculated from the technical replicates. An RT–qPCR assay for Actin Beta (ActB) transcripts was used as a housekeeping transcript reference to calculate $\Delta C_t$ values. Fold change values were calculated with the $\Delta\Delta C_t$ method using the average $\Delta C_t$ of all controls (DMSO-treated cells) as a reference.

#### RNAseq of MOLM-13 cells

RNAseq libraries were prepared using 100 ng of total RNA per sample with the NEBnext Ultra II Directional RNA library kit for Illumina (New England Biolabs) according to the manufacturer's recommendations. Libraries were quantified with the NEBnext library quant kit for Illumina (NEB) and then sequenced for 100 nt using a V3 single read flow cell on a HiSeq 2500 (Illumina) to an average read depth of 36 M reads per sample. Exact numbers for each sample used in this study are listed in the Supplementary Information. After QC with FastQC Version 0.11.2 (http://www.bioin forma tics.bbsrc.ac.uk/proje cts/fastq c/), reads were adaptor-trimmed (Q < 20) with Cutadapt (Version 1.4.132) using Trim Galore! (Version 0.3.3) (https://www.bioin forma tics.babra ham.ac.uk/proje cts/trim_galor e/). Reads were aligned to hg38 assembly with the grape-nf pipeline (https://github.com/guigo lab/grape -nf) wrapping STAR [5] (Version 2.4.0j33) and RSEM [6] (Version 1.2.2134). Differential analysis was performed in R using the DESeq2 package [7].

The sample reads (M) for each MOLM-13 RNAseq sample presented in this study were the following:

- DMSO Ctrl.: 1 – 40.47, 2 – 23.72, 3 – 32.16
- cAzadC (0.5 µM, 72 h): 1 – 32.34, 2 – 17.53, 3 – 4.83

#### LC-MS/QQQ analysis

For quantitative mass spectrometry an Agilent 1290 Infinity equipped with a variable wavelength detector (VWD) combined with an Agilent Technologies G6490 Triple Quad LC/MS system with electrospray ionization (ESI-MS, Agilent Jetstream) was used. Operating parameters: positive-ion mode, cell accelerator voltage of 5 V, $N_2$ gas temperature of 120 °C and $N_2$ gas flow of 11 L/min, sheath gas ($N_2$) temperature of 280 °C with a flow of 11 L/min, capillary voltage of 3000 V, nozzle voltage of 0 V, nebulizer at 60 psi, high-pressure RF at 100 V and low-pressure RF at 60 V. The instrument was operated in dynamic MRM mode (Table S10). For separation an Uptisphere C18-HDO column (3.0 µm, 150 × 2.1 mm from Interchim, UP3HDO-150/021) was used. Running conditions were 35 °C and a flow rate of 0.35 mL/min in combination with a binary mobile phase of 5 mm NH4OAc aqueous buffer A, brought to pH 4.9 with glacial acetic acid (200 µL/L), and an organic buffer B of 2 mm $NH_4COOH$ in 80 % acetonitrile (Roth, Ultra LC-MS grade, purity ≥ 99.98). The gradient started at 100 % solvent A for 0.5 min, followed by an increase of solvent B to 10 % over 5.5 min. From 6.0 min to 8.5 min, solvent B increased to 20 % then to 80 % in 1 min and maintained at 80 % for 1.5 min before returning to 100 % solvent A in 0.5 min and a 2.2 min re-equilibration period. Of each sample 10 µL were co-injected with 1 µL of stable isotope labeled internal standard (ISTD) which was aspirated automatically before each injection from the instrument itself. The sample data were analyzed by the quantitative and qualitative MassHunter

Software from Agilent using the integrated calibration function. The calibration solutions ranged from 0.1 pmol to 200 pmol for each canonical nucleoside and from 0.004 pmol to 5 pmol for each modified nucleoside (12 calibration levels).

α-Ketoglutarate (αKG) assay

To determine the relative amount of a-KG, MOLM-13 cells were seeded at a concentration of 5.0 x $10^5$ cells/mL and treated for 72 h with 0.5 µm cAzadC or AzadC, respectively. Samples (n = 8 – 9 biologically independent replicates) were analyzed using the α-Ketoglutarate Assay Kit (Sigma-Aldrich #MAK054) according to the manufacturer's protocol. After harvest, the cells were washed twice with PBS and resuspended in 100 µL α-KG buffer. Samples were filtered for 15 min at 13000 *x g*, 4°C using Ultrafree Centrifugal Filter Units (Sigma-Aldrich #UFC30LG25). The flow through was collected and divided into an αKG assay sample and an αKG assay blank well to have an individual blank value for each sample. The Fluorescent Peroxidase Substrate was diluted 5-fold in α-KG buffer prior to use. All samples were measured in a final volume of 100 µl of α-KG buffer supplemented with 2 µL αKG Converting Enzyme, 2 µL αKG Development Enzyme and 2 µL Fluorescent Peroxidase Substrate per sample. All blanks were measured in a final volume of 100 µL of αKG buffer containing 2 µL αKG Development Enzyme and 2 µL Fluorescent Peroxidase Substrate per sample. Samples were measured as a fluorometric assay at $\lambda_{ex}$ = 535 nm /$\lambda_{em}$ = 587 nm on a multimode microplate reader (Tecan). The normalized relative fluorescent unit (RFU), which correlates with the amount of αKG, was calculated for each sample by subtracting the RFU of the blank from the RFU of the sample.

Flow-cytometry based proliferation and apoptosis assays

The flow-cytometry based proliferation and apoptosis assays in MOLM-13 cells were performed as previously described.[8]

*Ex vivo* studies with AML-491 PDX cells

AML-491 PDX cells were re-isolated from full-blown donor mice and cultivated in StemPro-34 medium (Thermo Fisher Scientific Waltham, MA, USA) supplemented with 1 % l-Glutamine, 1 % Penicillin/Streptomycin, 2 % FCS (all Gibco, San Diego, CA, USA), 10 ng/mL hrFLT3L (R&D Systems, Minneapolis, MN, USA), 10 ng/mL hrSCF, 10 ng/mL hrTPO, and 10 ng/mL hrIL3 (all Peprotech, Rocky Hill, NJ, USA) at a density of 1 x $10^6$ cells per mL, and kept at 37 °C, 5% $CO_2$. Cells were treated with 0.5 µM, 1.0 µM or 3 µM cAzadC or AzadC for 72 h, then washed once in PBS, and either stored at -80 °C until further sample preparation for QQQ-MS analysis or RT-qPCR.

*In vivo* therapy trials

9 to 16 weeks old male NOD.Cg-Prkdc$^{scid}$ IL2rg$^{tm1Wjl}$/SzJ (NSG) mice (Charles River, Sulzfeld, Germany) were transplanted into the tail vein with 1.0 – 1.5 × $10^6$ luciferase-positive AML-491 PDX cells. Tumor outgrowth was monitored by *in vivo* bioluminescence imaging (BLI) [9]. When total flux reached values around 1 × $10^9$ Photons / second (around d21 after injection), mice were treated as indicated (cAzadC and AzadC were diluted in PBS) by intraperitoneal injection once

per day, 5 days a week for four consecutive weeks. Leukemic burden was monitored by BLI every 1 – 4 weeks. Body weight and general health condition were checked daily. Mice were sacrificed at advanced leukemic disease (BLI signal >$10^{10}$ Photons/sec), when clinical signs of illness appeared (rough fur, hunchback, reduced motility), if therapy related toxicity was observed (loss of body weight >15 %), or if BLI was below detection limit for more than three months.

**References Supplementary Information**
